# Functional characterization of *Botrytis cinerea* ABC transporter gene *BcatrB* in response to phytoalexins produced in plants belonging to families Solanaceae, Brassicaceae and Fabaceae

**DOI:** 10.1101/2022.11.07.515369

**Authors:** Abriel Salaria Bulasag, Maurizio Camagna, Teruhiko Kuroyanagi, Akira Ashida, Kento Ito, Aiko Tanaka, Ikuo Sato, Sotaro Chiba, Makoto Ojika, Daigo Takemoto

**Author notes:** **Corresponding Author:** Daigo Takemoto.

## Abstract

*Botrytis cinerea*, a generalist fungal pathogen of economically important crop species, has been shown to exhibit reduced sensitivity to fungicides and plant toxins. Specifically, previous reports indicate *B. cinerea*’s efficacy in tolerating a wide array of phytoalexins, toxic plant metabolites that play key role in plant immune defense strategies. Previously, we have shown that a distinct set of genes was induced in *B. cinerea* when treated with phytoalexins derived from different plant species such as rishitin (tomato and potato), capsidiol (tobacco and bell pepper) or resveratrol (grape and blueberry). In this study, we focused on the functional analyses of *B. cinerea* genes induced by rishitin treatment. *B. cinerea* can metabolize rishitin to at least 4 oxidized forms. Heterologous expression of rishitin-induced *B. cinerea* genes in the plant symbiotic fungus, *Epichloë festucae*, revealed that oxidoreductase (Bcin08g04910) and cytochrome P450 (Bcin16g01490) genes are involved in the oxidation of rishitin. BcatrB is an exporter of structurally unrelated anti-microbial compounds such as resveratrol, camalexin and fungicide fenpicionil. Expression of *BcatrB* is upregulated by rishitin, but not by structurally resembling capsidiol. *BcatrB* knock out transformants (*ΔbcatrB*) showed enhanced sensitivity to rishitin, but not to capsidiol. Likewise, *ΔbcatrB* showed reduced virulence on tomato fruits (which produce rishitin), but showed full virulence on *Nicotiana benthamiana* (which mainly produces capsidiol), suggesting that *B. cinerea* distinguishes phytoalexins and activates expression of appropriate transporter genes during the infection. Activation of *BcatrB* promoter was detected using P_*BcatrB*:*GFP* transformant during the *B. cinerea* infection in plant tissues. Surveying of 26 plant species across 13 families revealed that the *BcatrB* promoter is mainly activated during the infection of plants belonging to the Solanaceae, Fabaceae and Brassicaceae families. The *BcatrB* promoter is activated by the treatment with Fabaceae phytoalexins medicarpin and glyceollin, and *ΔbcatrB* showed reduced virulence on red clover, which produces medicarpin. These results suggest that *BcatrB* plays a critical role in the strategy employed by *B. cinerea* to bypass the plant innate immune responses in a wide variety of important crops belonging to the Solanaceae, Brassicaceae and Fabaceae families.

## INTRODUCTION

*Botrytis cinerea*, commonly known as grey mold, is one of the most economically important pathogens. It affects approximately 1,400 plant species across several plant families (Elad, 2016) and brings severe yield losses to high impact crops such as grapevine, Solanaceae (tomato, potato, bell pepper), Brassicaceae (canola), Fabaceae (pea, bean), and among others. However, recent studies suggested that its potency as a pathogen is much more complex than previously described (Frías et al., 2016). It is generally held that *B. cinerea* craftily manipulates plant defenses, resulting in plant cell death by means of plant cell death inducing proteins (PCIDs) (Petrasch et al., 2019; Leisen et al., 2022), thereby creating a favorable environment for necrotrophy. Likewise, it stimulates the plant host to an exchange of enzymes and toxins, promoting disease progression (Bi et al., 2022). The evolutionary path taken by these polyxenous pathogens to diversify their host range is particularly intriguing, but molecular mechanisms driving these evolutionary changes are largely unknown. Several studies suggest that the virulence across plant species is enabled by complex interactions that include key fungal processes such as oxidative stress response (N’Guyen et al., 2021) and cell wall integrity (Cui et al., 2021; Escobar-Niño et al., 2021).

As a necrotrophic pathogen, *B. cinerea* can infect a wide variety of plant hosts by dexterously bypassing various aspects of a plant’s immune arsenal. One such critical component in plants, shaped and diversified through evolutionary lineages and plant-pathogen interaction events, are a group termed as phytoalexins. Fungal pathogens must be able to nullify these potent toxins to launch a successful attack. This presupposes the development of complex and coordinated toolkits to metabolize an equally diverse plant arsenal to launch an effective counterattack. Such plant antimicrobials were also deemed crucial to the development of non-host resistance to necrotrophs (Sanchez-Vallet et al., 2010), inadvertently prompting fungal pathogens to broaden their host repertoire.

Another significant hurdle confronting pathogens is that several of these plant chemicals act in unison and occur in such diversity to challenge such dexterous pathogens (Pedras and Abdoli, 2017; Allan et al., 2019; Newman et al., 2020; N’Guyen et al., 2021; Westrick et al., 2021). Several studies attribute this dexterity to detoxification of phytochemicals to multi-functional enzymes such as cytochrome P450 (oxidoreductase, monooxygenase and dehydratases) which are quintessential components of robust fungal pathogenic systems (Pedras and Ahiahonu, 2005; Kuroyanagi et al., 2022).

Moreover, some generalist pathogens employ an array of molecular transporters to evade each plant host’s chemical weaponry. This offers another layer of host-pathogen interaction that could explain host range expansion. Three fundamental selection factors: alternative nutrient sources, niche range expansion and establishment in unconventional environmental contexts as key drivers to the evolution of fungal transporters (Milner et al., 2019). Fungal transporters, encoded by singular genes, can be conveniently transferred across fungal phyla through horizontal gene transfer, leading to enhanced polyxenous behavior. Using combined transcriptomic and metabolomic approaches, it was found that multidrug resistance to fungicides is primarily aided by efflux transporter genes enhancing virulence of key fungal pathogens (Zhang et al., 2019; Samaras et al., 2020; Shi et al., 2020; Wang et al., 2021; Bartholomew et al., 2022; Harper et al., 2022; Wang et al., 2022). More specifically, ABC (ATP-binding cassette) and MFS (major facilitator superfamily) transporters dispose key phytoalexins by exporting them out of fungal cells (Newman et al., 2020; Kumari et al., 2021; Madloo et al., 2021; Westrick et al., 2021) paving the way to generalist pathogen lineages.

In summary, the mechanism conferring generalist pathogens to drastically expand their host range is likely driven by two important processes: 1) enzymatic detoxification of key antimicrobial metabolites; and 2) efflux processes facilitated by membrane transporters. Hence, this study aspires to provide a novel perspective in understanding molecular mechanisms governing the interaction between phytoalexin production in various plant model systems and its subsequent metabolism and efflux in the polyxenous grey mold pathogen *B. cinerea*.

## MATERIALS AND METHODS

### Biological Material, Growth Conditions and Incubation in Phytoalexins

*B. cinerea* strain AI18 (Kuroyanagi et al., 2022), *Epichloë festucae* strain Fl1 and their transformants used in this study are listed in Supplementary Table S1. They were grown on potato dextrose agar (PDA) at 23°C. For the incubation in phytoalexins, mycelia plugs (approx.1 mm^3^) were excised from the growing edge of the colony using a dissection microscope (Stemi DV4 Stereo Microscope, Carl Zeiss, Oberkochen, Germany) and submerged in 50 μl of water or indicated phytoalexin in a sealed 96 well clear plate. The plate was incubated at 23°C for the indicated time.

Capsidiol was purified from *Nicotiana tabacum* as previously reported (Matsukawa et al., 2013) and synthesized rishitin (Murai et al., 1975) was provided by former Prof. Akira Murai (Hokkaido University, Japan). Resveratrol and brassinin were obtained from Sigma-Aldrich (Burlington, MA, USA). Glyceollin (glyceollin I) was obtained from Wako pure chemical (Osaka, Japan). Medicarpin was obtained from MedChemExpress (Monmouth Junction, NJ, USA).

### Detection of Rishitin and Their Metabolites Using LC/MS

For the detection of rishitin and their metabolites after the incubation with *B. cinerea* or *E. festucae* transformants, the supernatant (50 μl) was collected, mixed with 50 μl acetonitrile and measured by LC/MS (Accurate-Mass Q-TOF LC/MS 6520, Agilent Technologies, Santa Clara, CA, USA) with ODS column Cadenza CD-C18, 75 × 2 mm (Imtakt, Kyoto, Japan) as previously described (Kuroyanagi et al., 2022).

### RNAseq Analysis

Extraction of RNA and RNA sequencing analysis were performed as previously described (Kuroyanagi et al., 2022). The nucleotides of each read with less than 13 quality value were masked and reads shorter than 50 bp in length were discarded, and filtered reads were mapped to annotated cDNA sequences for *B. cinerea* (ASM83294v1, GenBank accession GCA_000143535) using Bowtie software (Langmead et al., 2009). For each gene, the relative fragments per kilobase of transcript per million mapped reads (FPKM) values were calculated and significant difference from the control was assessed by the two-tailed Student’s *t*-test. RNA-seq data reported in this work are available in GenBank under the accession numbers DRA013980.

### Extraction of Genomic DNA, PCR and Construction of Vectors

Genomic DNA of *E. festucae* and *B. cinerea* was isolated from fungal mycelium grown in potato dextrose broth (PDB) using DNeasy Plant Mini Kit (QIAGEN, Hilden, Germany). PCR amplification from genomic and plasmid DNA templates was performed using PrimeStar Max DNA polymerase (Takara Bio, Kusatsu, Japan) or GoTaq Master Mix (Promega, Madison, WI, USA). Vectors for heterologous expression, detection of promoter activity, gene knock out used in this study are listed in Supplementary Table S2. Sequences of primers used for the construction of vectors and PCR to confirm the gene knockout are listed in Supplementary Table S3.

### Fungal Transformation

Preparation of *E. festucae* and *B. cinerea* protoplasts and transformation was performed as previously reported (Kuroyanagi et al., 2022). Candidate colonies were exposed to BLB blacklight for the induction of sporulation and single spore isolation was performed to obtain purified strains. Note that *ΔbcatrB*-14 and -23 were isolated from separate transformation experiments. Transformants of *E. festucae* and *B. cinerea* used in this study are listed in Supplementary Table S1.

### Pathogen Inoculation

Leaves or fruits (tomato, grape) of plant species were kept in moistened and sealed in a plastic chamber. Leaves detached from the plant were covered with a wet Kimwipes at the cut end of the stem. Mycelial plugs (approx. 5 mm x 5 mm) of *B. cinerea* were excised from the growing edge of the colony grown on PDA and placed on the abaxial side of the leaf or on the fruit and covered with wet lens paper. For the inoculation on tomato, mycelial blocks of *B. cinerea* were placed on the cut surface of tomato, and the fruits were kept at high humidity at 23°C for 5 days. *B. cinerea* conidia formed on tomato were washed off in 15 ml water, and number of conidia in water was counted using hemocytometer.

### Microscopy

Images of *B. cinerea* expressing GFP under the control of the *BcatrB* promoter were collected using a confocal laser scanning microscope FV1000-D (Olympus, Tokyo, Japan). The laser for detection of GFP was used as the excitation source at 488 nm, and GFP fluorescence was recorded between 515 and 545 nm. Images were acquired with settings that did not saturate the fluorescence, and the total fluorescence per spore was determined using ImageJ software (Schneider et al., 2012).

### Detection of Luciferase Activity in *B. cinerea* P_*BcatrB*:*Luc* Transformant

*B. cinerea* P_*BcatrB*:*Luc* transformant was grown on PDA at 23°C. Three mycelia plugs (approx. 2 mm x 2 mm) were excised from the growing edge of the colony and submerged in 50 μl of water or indicated phytoalexin containing 50 μM D-luciferin in a sealed 96-well microplate (Nunc 96F microwell white polystyrene plate, Thermo Fisher Scientific, Waltham, MA, USA). Changes in luminescence intensity were measured over time with Mithras LB 940 (Berthold Technologies, Bad Wildbad, Germany).

### Bioinformatics

Sequence data was analyzed and annotated using MacVector (version 18.2 or earlier; MacVector Inc., Apec, NC, USA).

## RESULTS AND DISCUSSION

### Rishitin Treatment Induced Genes Predicted to be Involved in the Metabolization and Efflux of the Phytoalexin in *B. cinerea*

Previously, we have performed RNA-seq analysis of *B. cinerea* in response to sesquiterpenoid (rishitin and capsidiol) and stilbenoid (resveratrol) phytoalexins. *Bccpdh*, encoding a short chain dehydrogenase, was identified as a gene specifically induced in *B. cinerea* treated with capsidiol, and BcCPDH was revealed to be involved in the detoxification of capsidiol to less toxic capsenone (Kuroyanagi et al., 2022). In this study, we focused on genes induced by rishitin treatment. Further analysis of the RNAseq data identified rishitin-induced genes belonging to various groups, such as cytochrome P450/oxidoreductases, ABC transporters, cell wall degrading enzymes (CWDEs), and secondary metabolite synthesis (Figure 1). Some genes were identified to be potentially involved in the metabolization of rishitin, to convert it into its oxidized form, including an oxidoreductase (Bcin08g04910) and cytochrome P450 genes (Bcin16g01490, Bcin06g00650, Bcin07g05430) (Figure 1A, Kuroyanagi et al., 2022). As such these genes were investigated further in terms of their ability to metabolize rishitin (see next section). Expression of several ABC transporter genes was also induced, and may thus be involved in the efflux of rishitin to enhance the tolerance of *B. cinerea* (Figure 1B). ABC transporter BcatrB has been reported as an exporter of structurally unrelated phytoalexins such as resveratrol, camalexin and fungicides (Schoonbeek et al., 2001; Vermeulen et al., 2001; Stefanato et al., 2009). Our RNAseq data indicated that expression of *BcatrB* was induced in rishitin and resveratrol, but not in capsidiol. Conversely, expression of *BcatrD* and *Bmr3* was significantly higher during rishitin treatment as compared to other phytoalexins. These genes could be responsible for efflux of rishitin to enhance the tolerance of *B. cinerea*. A number of studies link the transcriptional activation of efflux ABC transporters such as *BcatrB* and *BcatrD* to induction by phytoalexins and fungicides (Hayashi et al., 2001; Perlin et al., 2014). On the other hand, upregulation of *Bmr1* and *Bmr3* has been reported in response to several fungicides, a phytoalexin (resveratrol) and toxicants (Makizumi et al., 2002). While the four ABC transporter genes described above are induced in rishitin-treated *B. cinerea*, expression of a different set of transporter genes is activated when treated with capsidiol (Figure 1B, Kuroyanagi et al., 2022). These results suggest that the effective transporters differ depending on the phytoalexins, and *B. cinerea* could be inducing expression of the appropriate transporter genes by recognizing different phytoalexins. In the later part of this manuscript, we focused on the functional analysis of *BcatrB*, especially its contribution to the tolerance of *B. cinerea* to rishitin.

Interestingly, some genes that are not directly involved in the detoxification of phytoalexins, but instead related to the virulence of *B. cinerea*, are upregulated in *B. cinerea* treated with rishitin. For example, expression of genes encoding CWDEs, such as genes for polygalacturonase *Bcpg1* (Bcin14g00850), xylanase *BcXyn11a* (Bcin02g01960) and glycosyl hydrolase (Bcin02g01960), were induced by rishitin treatment (Figure 1C). Previous studies have elaborated on the role of CWDEs as virulence factors in *B. cinerea*. Knockout of a polygalacturonase gene *Bcpg1* had no effect on primary infection, but caused significant decrease in secondary infection (development of the lesion) (ten Have et al., 1998). *BcXyn11a*, encoding an endo-*β*-1,4-xylanase for the degradation of hemicellulose, is shown to be required for full virulence of *B. cinerea* (Brito et al., 2006). Genes that are part of a subtelomeric cluster for the production of phytotoxic botcinic acid are also activated by rishitin and capsidiol treatment (Figure 1D). Given that these genes involved in the spread of disease symptoms are also induced by rishitin in *B. cinerea*, implies that phytoalexins produced by plants as resistance factors are used by *B. cinerea* as cues to promote virulence.

**FIGURE 1.**
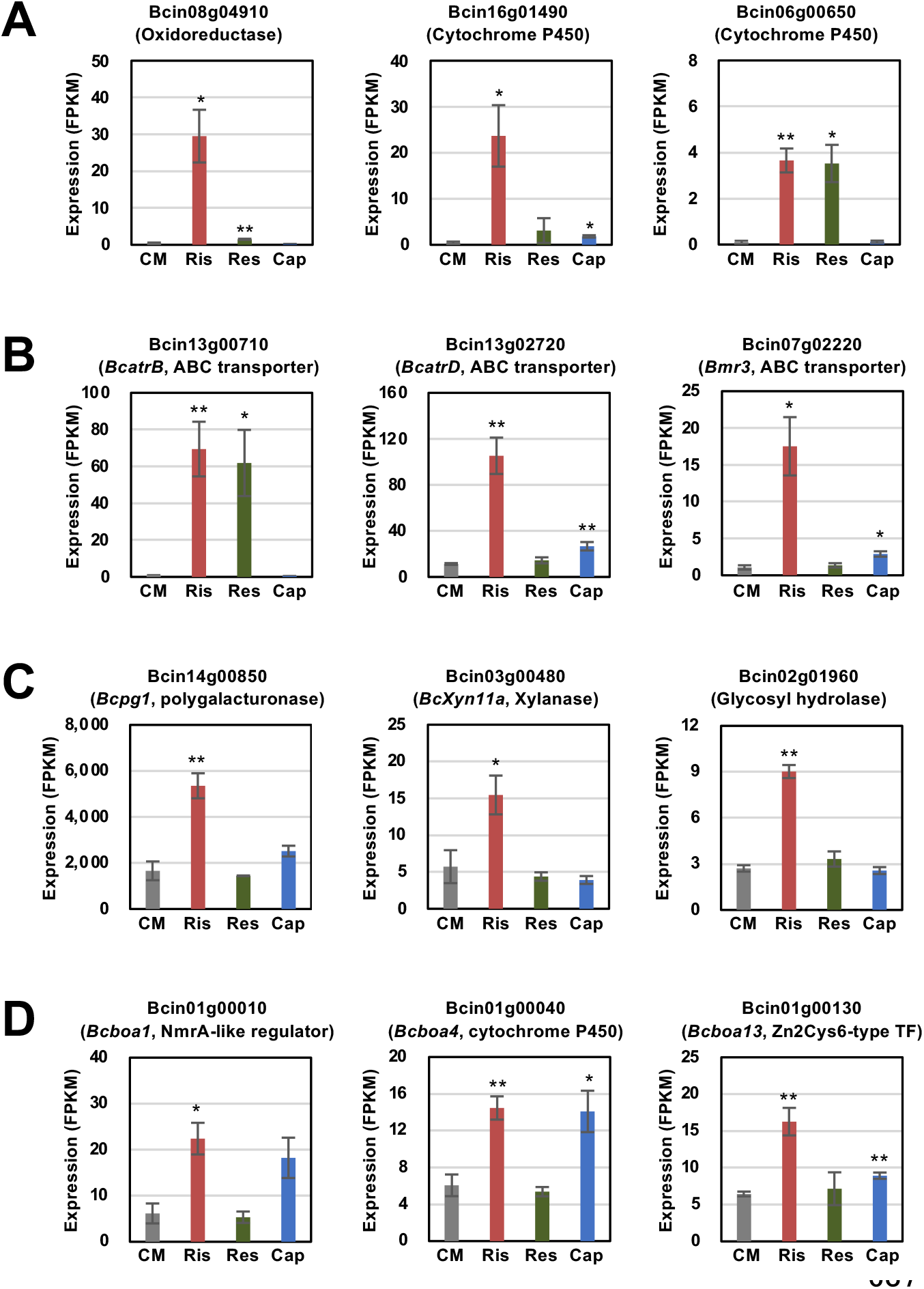
Transcriptomic changes in *Botrytis cinerea* genes induced by rishitin. **(A)** Cytochrome P450/oxidoreductases **(B)** ABC transporters **(C)** Cell wall degrading enzymes (CWDEs) and **(D)** Botcinic acid biosynthesis. TF, transcription factor. The gene expression (FPKM value) was determined by RNA-seq analysis of *B. cinerea* cultured in CM media containing 500 µM rishitin, 500 µM resveratrol or 100 µM capsidiol for 24 h. Data are mean ± SE (*n* = 3). Asterisks indicate a significant difference from the control (CM) as assessed by two-tailed Student’s *t*-test, \*\**P* < 0.01, \**P* < 0.05.

### Heterologous Expression of Rishitin-induced *B. cinerea* Genes in Symbiotic Fungus Results in the Metabolization of Rishitin to its Oxidized Forms

Rishitin is metabolized to at least 4 oxidized forms by *B. cinerea* (Figure 2A, Kuroyanagi et al., 2022). To investigate the function of *B. cinerea* cytochrome P450 and oxidoreductase genes that were significantly upregulated in rishitin treatment (Figure 1A), candidate genes were selected to be heterologously expressed in grass symbiotic fungus *Epichloë festucae* to detect the enzymatic activity of the encoded proteins. Kuroyanagi et al. (2022) have used the same system to identify a dehydrogenase BcCPDH, which can convert capsidiol to capsenone. Two genes upregulated in *B. cinerea* treated with rishitin, Bc08g04910 and Bc16g01490, were expressed in *E. festucae* under the control of the TEF promoter (Vanden Wymelenberg et al., 1997) for constitutive expression. These *E. festucae* transformants were incubated in rishitin solution to examine their function in rishitin oxidation. Two days after incubation in 100 µM rishitin, the *E. festucae* strain expressing Bc08g04910 caused reduction of rishitin and oxidized rishitin was detected (Figure 2B). Similarly, oxidized rishitin was detected after the incubation of rishitin with the *E. festucae* strain expressing Bc16g01490, although a significant reduction of rishitin was not observed at 2 days. Reduction of rishitin and pronounced production of two oxidized rishitin derivatives was observed after 10 days of incubation with *E. festucae* expressing Bc16g01490 (Supplementary Figure 1). *E. festucae* control strains expressing *DsRed* gene didn’t induce a reduction or oxidation of rishitin (Figure 2B). While *B. cinerea* produced various rishitin metabolites after the incubation with rishitin, each of the *E. festucae* transformants showed one or two peaks which demonstrated a mass increase indicative of the presence of an additional oxygen atom in the rishitin molecule. These results suggests that oxidoreductases (Bc08g04910) and cytochrome P450 (Bc16g01490) can metabolize rishitin into oxidized forms, but there should be additional genes (enzymes) involved in the rishitin oxidation. Employing these two *E. festucae* transformants in large scale incubation with rishitin should enable further insights into the chemical structure of the oxidized rishitin compounds.

**FIGURE 2.**
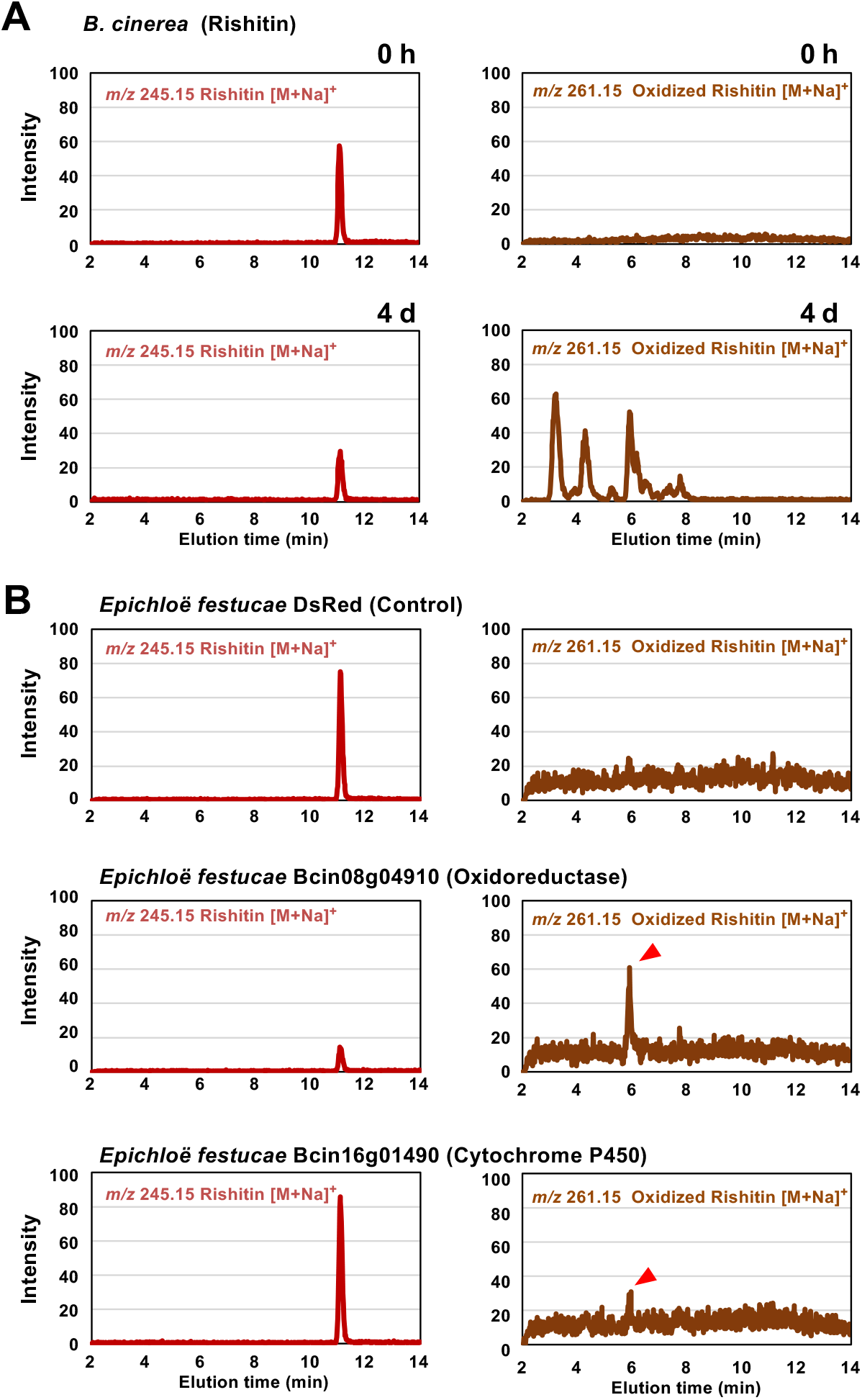
Metabolization of rishitin by *Botrytis cinerea* and *Epichloë festucae* transformants expressing rishitin-induced *B. cinerea* genes. **(A)** Mycelial block (approx. 1 mm^3^) of *B. cinerea* was incubated in 50 µl of 500 µM rishitin for 4 days and remaining rishitin and oxidized rishitin were detected by LC/MS. **(B)** Mycelial block (approx. 1 mm^3^) of *E. festucae* transformants expressing *DsRed* gene (control) or rishitin-induced *B. cinerea* genes (Bcin08g04910 or Bcin16g01490) were incubated in 50 µl of 100 µM rishitin for 2 days and remaining rishitin and oxidized rishitin were detected by LC/MS. See Supplementary Figure S1 for 10 days incubation of *E. festucae* transformants expressing Bcin16g01490 in rishitin.

This study has yet to identify all genes that correspond to the metabolization of rishitin into several oxidized forms by the wild-type *B. cinerea* strain. Multiple genes are involved in the detoxification/oxidation of rishitin (Figure 2B), KO of single gene may therefore not have a significant effect on the virulence of this pathogen. In some cases, metabolism for detoxification of phytoalexins may only involve the action of a single enzyme, such as in the case of capsidiol (Kuroyanagi et al., 2022) and pisatin (Matthews and Van Etten, 1983). On the other hand, a series of enzymatic reactions catalyze the conversion of cruciferous phytoalexins such as brassinin and camalexin into their less toxic forms (Pedras and Abdoli, 2017). This was shown across a range of fungal pathogens including necrotrophs such as *B. cinerea* and *Sclerotinia sclerotiorum*. However, enzymes catalyzing detoxification reactions in these pathogens are still yet to be determined. Despite similar structural profiles of capsidiol and rishitin, *B. cinerea* employs different mechanisms for a similar xenobiotic detoxification process.

### *BcatrB* KO Mutants Showed Increased Sensitivity to Rishitin

Previous studies on *BcatrB* established its role in the tolerance of *B. cinerea* to structurally unrelated phytoalexins such as resveratrol, camalexin and the fungicide fenpicionil (Schoonbeek et al., 2001; Vermeulen et al., 2001; Stefanato et al., 2009). Given that the expression of *BcatrB* is upregulated by rishitin (Figure 1B), *BcatrB* is expected to be also involved in the tolerance of *B. cinerea* to rishitin.

To investigate the role of *BcatrB* in rishitin tolerance of *B. cinerea, BcatrB* knock out strains were produced (Supplementary Figure 2). Conidial germination of wild type and *BcatrB* KO strains was measured after treatment with rishitin or capsidiol (Figure 3A). In 100 µM capsidiol, the length of *B. cinerea* germ tubes were shortened compared to those without treatment (Figure 3A). However, KO of *BcatrB* had no significant effect on the sensitivity of *B. cinerea* to capsidiol, which is consistent with the fact that *BcatrB* is not induced by capsidiol treatment (Figure 1B). In contrast, *BcatrB* mutants showed enhanced sensitivity to rishitin compared with wild type (Figure 3A).

**FIGURE 3.**
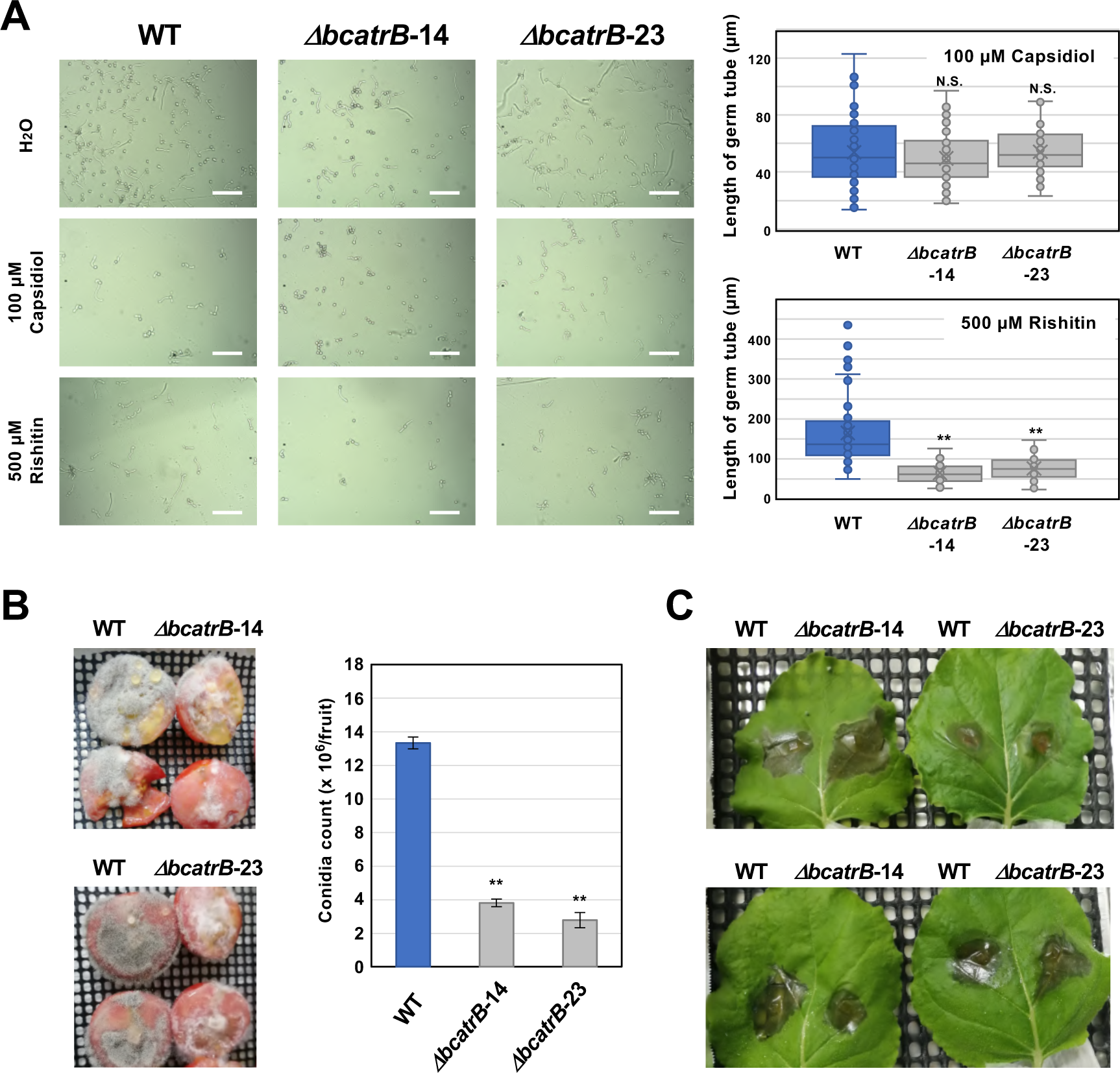
Deletion of *BcatrB* gene compromises the rishitin tolerance of *Botrytis cinerea*. **(A)** Conidial suspension of *B. cinerea* was incubated in water (H_2_O), 100 µM capsidiol or 500 µM rishitin and the length of germ tube was measured after 18 h incubation. Bars = 100 µm. Data are mean ± SE (n = 60). Asterisks indicate a significant difference from wild type (WT) as assessed by two-tailed Student’s *t*-test. \*\**P* < 0.01. N. S., not significant. Lines and crosses (x) in the columns indicate the median and mean values, respectively. **(B)** Tomato fruits (cut in half) were inoculated with mycelia plug (approx. 5 × 5 mm) of *B. cinerea* WT or *ΔbcatrB* strains and produced conidia were counted 5 days after the inoculation. Data are mean ± SE (n = 3). Asterisks indicate a significant difference from WT as assessed by two-tailed Student’s *t*-test. \*\**P* < 0.01. **(C)** Leaves of *Nicotiana benthamiana* were inoculated with mycelia plug (approx. 5 × 5 mm) of *B. cinerea* WT or *ΔbcatrB* strains. Photographs were taken 3 days after the inoculation.

Given that rishitin production has been detected in tomato fruits upon pathogen attack (Sato et al., 1968), virulence of *BcatrB* strains was tested on tomato fruits. Higher sporulation rate was observed for wild type as compared to *ΔBcatrB* strains in tomato fruits (Figure 3B). However, no significant differences between wild type and KO strains were observed in the lesion formation on *N. benthamiana* leaves (Figure 3C), which reportedly produce capsidiol as major phytoalexin (Shibata et al., 2016). These results suggest that *BcatrB* is involved in the tolerance of *B. cinerea* to rishitin, but not to capsidiol.

Phytoalexins induce damage to cell ultrastructure as well as conidial germination of *B. cinerea* (Adrian and Jeandet, 2012). While *BcatrB* has been shown to be an important efflux transporter for a variety of anti-microbial chemicals, other transporters have also been reported to be involved in resistance of *B. cinerea* to toxins. Deletion of MFS transporters in *B. cinerea, Bcmfsg* and *Bcmfs1*, resulted in the increase sensitivity to natural toxic compounds produced in plants (Hayashi et al., 2002; Vela-Corcía et al., 2019). These previous reports clearly demonstrate the use of multiple ABC and MFS transporters in reducing damage to *B. cinerea* by removal of these chemicals, natural or artificial, by an efflux transport mechanisms.

### *B. cinerea BcatrB* Promoter is Activated During the Infection in Plants Belonging to Solanaceae, Brassicaceae and Fabaceae

To search for the other phytoalexins that could be potential substrates of the BcatrB transporter, *B. cinerea* transformant expressing GFP under the control of the *BcatrB* promoter (P_*BcatrB_GFP*) were inoculated onto 25 host plants across 13 plant families (Figure 4, Supplementary Table S4).

**FIGURE 4.**
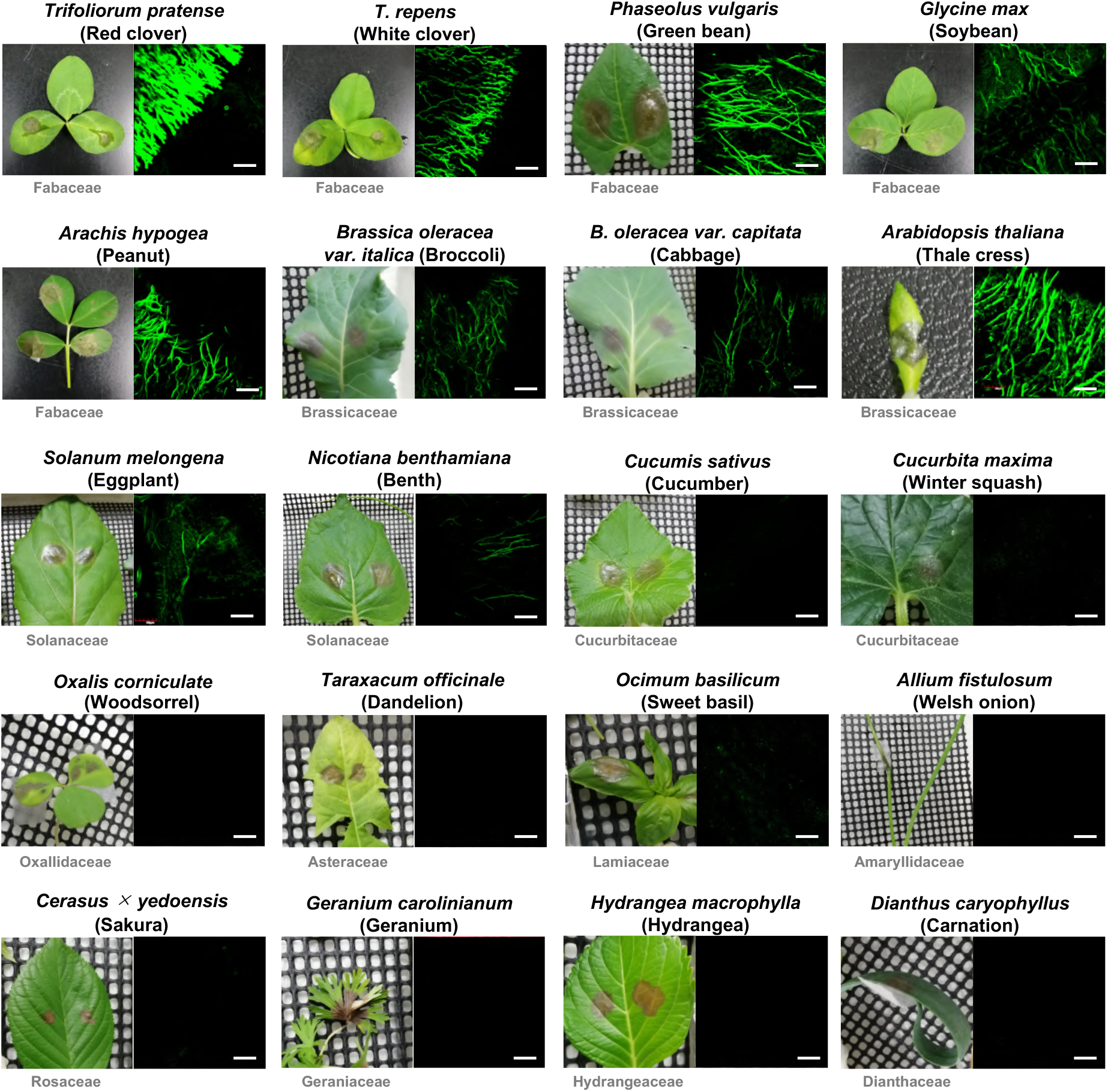
*Botrytis cinerea BcatrB* promoter is activated during the infection in Fabaceae, Brassicaceae and Solanaceae species. Leaves of indicated plants were inoculated with the mycelia of *B. cinerea* P_*BcatrB*:*GFP* transformant and hyphae at the edge of the lesion were observed by confocal laser microscopy 2 or 3 d after the inoculation. Bars = 100 µm.

Among the tested plant species, expression of GFP was detected mostly during the infection in Brassicaceae, Fabaceae and Solanaceae species (Figure 4). GFP expression ranged from weak, such as in the case of eggplant and *N. benthamiana*, to intense signals detected from red clover and Arabidopsis. Similarly, activity of the *BcatrB* promoter was perceived to widely deviate across members of the same family. The intensity of promoter activation also varied among plants at lower taxonomic levels such as in the case of red clover and white clover. Moreover, tissue-specific expression can also be observed for tomato and grape, with *BcatrB* expressed in fruits but weak or no expression in leaves (Figure 5). Interestingly, *BcatrB* expression can also be detected in infection cushions, which were only formed in fruit tissues (Figure 5). Differential expression of *BcatrB* might have been prompted by a higher production of rishitin and resveratrol in tomato and grape fruits, respectively. Infection cushions have been revealed to be formed alternatively with appressoria during less favorable conditions, to facilitate infection by penetration of the host tissue and upscaling virulence factors (Choquer et al., 2021, Bi et al., 2022).

**FIGURE 5.**
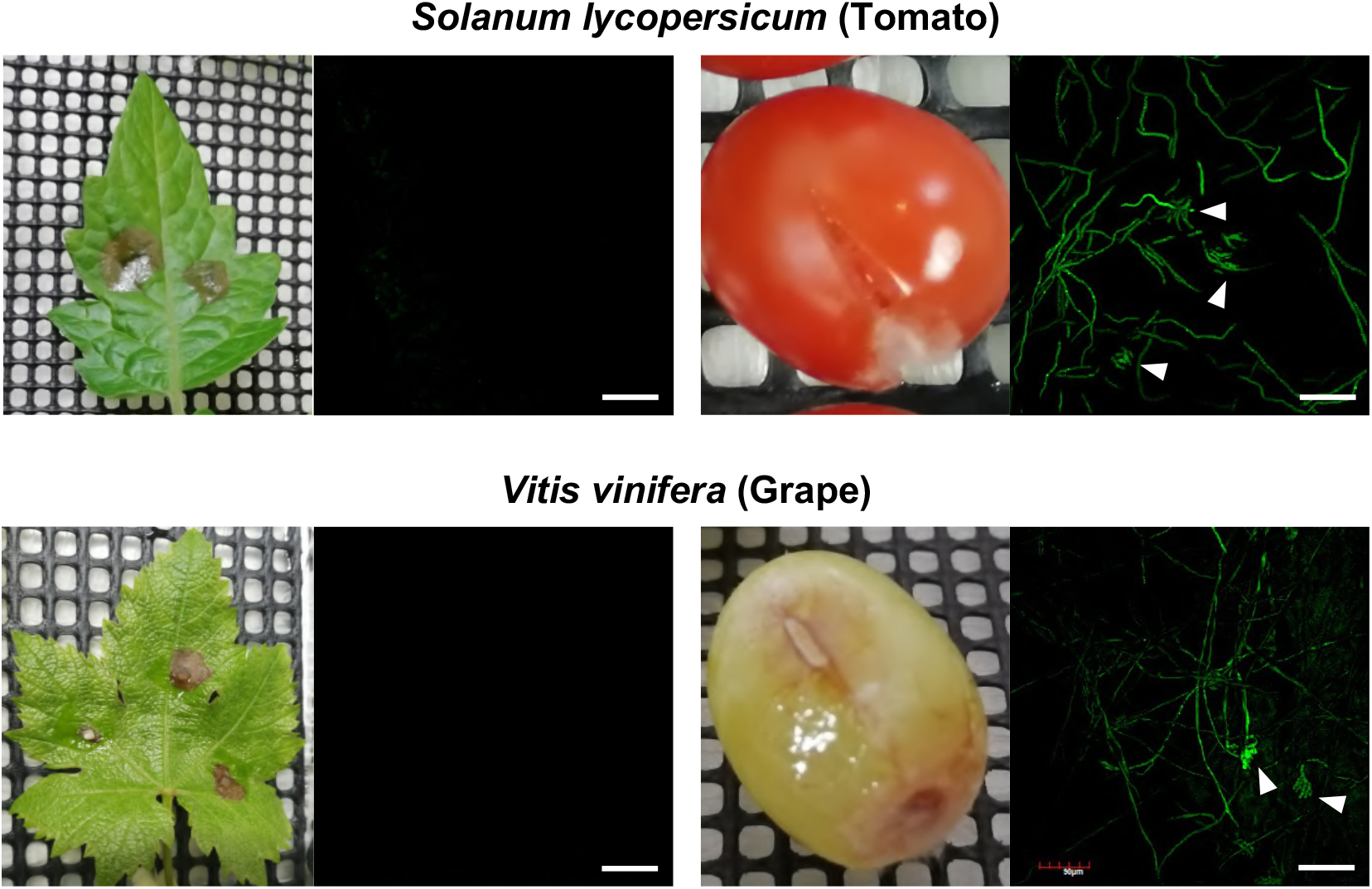
*Botrytis cinerea BcatrB* promoter is activated during the infection in fruits of tomato and grape. Leaves or fruits of tomato (top) or grape (bottom) were inoculated with the mycelia of *B. cinerea* P_*BcatrB*:*GFP* transformant and hyphae at the edge of the lesion was observed by confocal laser microscopy 2 or 3 d after the inoculation. Arrowheads indicate infection cushions. Bars = 100 μm.

### *ΔBcatrB* is Involved in the Virulence on Red Clover, a Fabaceae Plant Producing Pterocarpan Phytoalexins

Conidia of P_*BcatrB_GFP* transformants were treated with several phytoalexins from Brassicaceae, Fabaceae and Solanaceae species. Consistent with the RNAseq analysis data, the *BcatrB* promoter was activated by rishitin, but not by capsidiol (Figure 6A). The activation of the *BcatrB* promoter in *N. benthamiana* (Figure 4), which mainly produces capsidiol, is probably due to its response to other antimicrobial substances produced in *N. benthamiana*. Indole phytoalexins from Brassicaceae species, brassinin and camalexin, significantly activated the expression of GFP under the control of the *BcatrB* promoter. Pterocarpan phytoalexins produced in Fabaceae, medicarpin and glyceollin, also induced the activation of the *BcatrB* promoter (Figure 6A). Therefore, activation of the *BcatrB* promoter during the infection in Brassicaceae and Fabaceae plants (Figure 4) is presumed to occur via recognition of indole and pterocarpan phytoalexins by *B. cinerea*.

**FIGURE 6.**
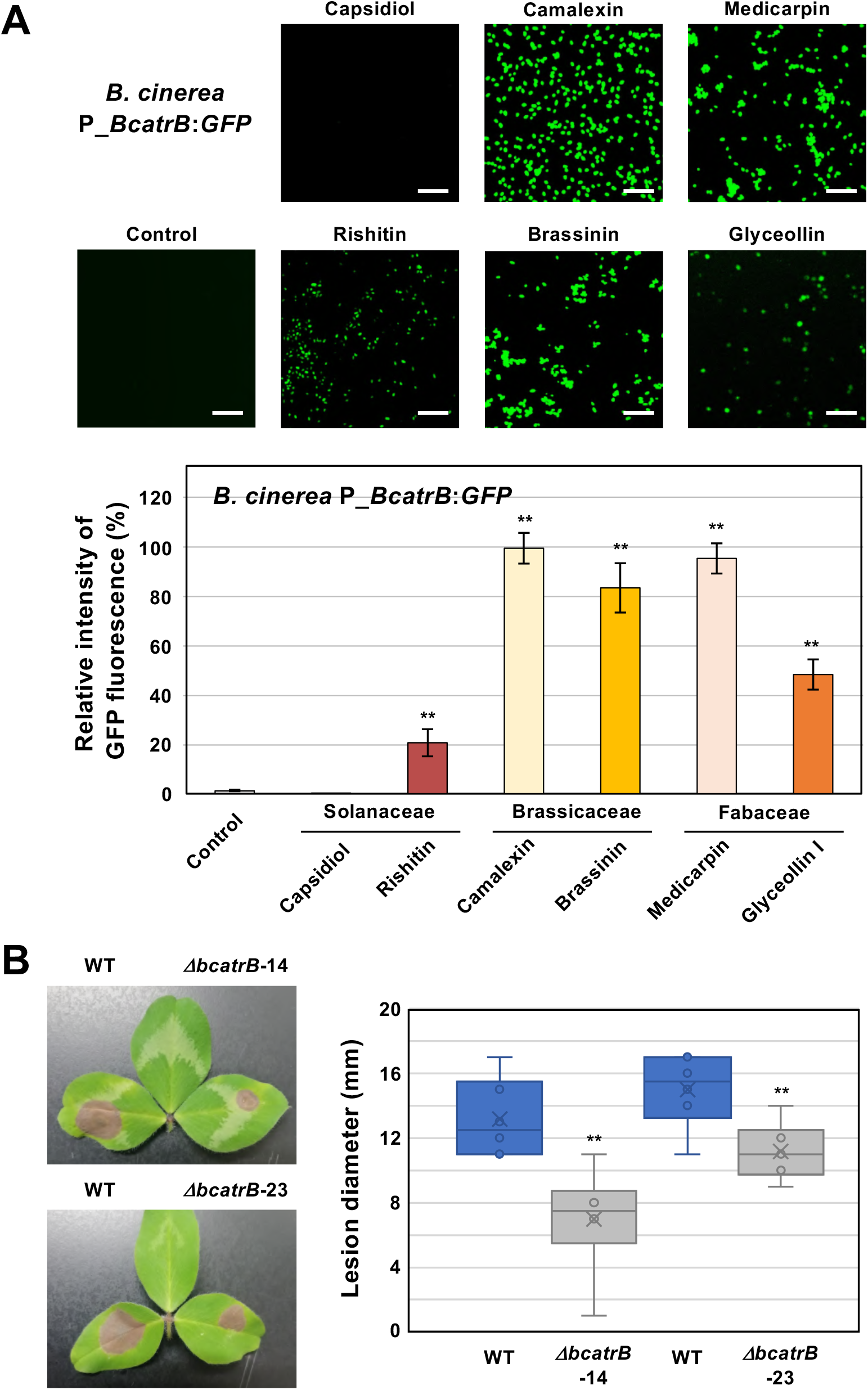
**(A)** *Botrytis cinerea BcatrB* promoter is activated by Solanaceae, Brassicaceae and Fabaceae phytoalexins. Conidia of *B. cinerea* P_*BcatrB*:*GFP* were treated with 1% DMSO (Control) or 100 µM of indicated phytoalexins, and GFP fluorescence was detected by confocal laser microscopy 2 d after the treatment. Bars = 100 µm. Data are mean ± SE (n = 20). Asterisks indicate a significant difference from control as assessed by two-tailed Student’s *t*-test. \*\**P* < 0.01. **(B)** Leaves of red clover (*Trifolium pratense*) were inoculated with mycelia plug (approx. 5 × 5 mm) of *B. cinerea* wild type (WT) or *ΔbcatrB* strains and lesion diameter was measured 3 days after the inoculation. Data are mean ± SE (n = 6). Asterisks indicate a significant difference from WT as assessed by two-tailed Student’s *t*-test. \*\**P* < 0.01.

Red clover (*Trifolium pratense*) had shown the most substantial activation of the *BcatrB* promoter (Figure 4). To further substantiate that *BcatrB* acts as a critical virulence factor for the infection of red clover, we performed inoculations using the *ΔBcatrB* KO strains. Compared to wild type inoculations, the *ΔBcatrB* strains resulted in lower lesion scores (Figure 6B), indicating that *BcatrB* is required for full virulence on red clover, which produces the pterocapan phytoalexins medicarpin and maackiain (Dewick 1975).

### *B. cinerea* Rapidly Activates the *BcatrB* Promoter in Response to Solanaceae, Brassicaceae and Fabaceae Phytoalexins

To investigate the activation profile of the *BcatrB* promoter in response to different phytoalexins, we produced a P_*BcatrB*:*Luc* transformant of *B. cinerea* which expresses a luciferase gene under the control of the 2 kb long promoter region of *BcatrB*. Activation of the *BcatrB* promoter was detected as luciferase-mediated chemiluminescence within 10 minutes after 100 µM rishitin treatment. Activation of the promoter reached its peak within 1 h and quickly declined (Figure 7). Kuroyanagi et al. (2022) reported that rishitin is completely metabolized into oxidized forms within 6 h after treatment initiation.

**FIGURE 7.**
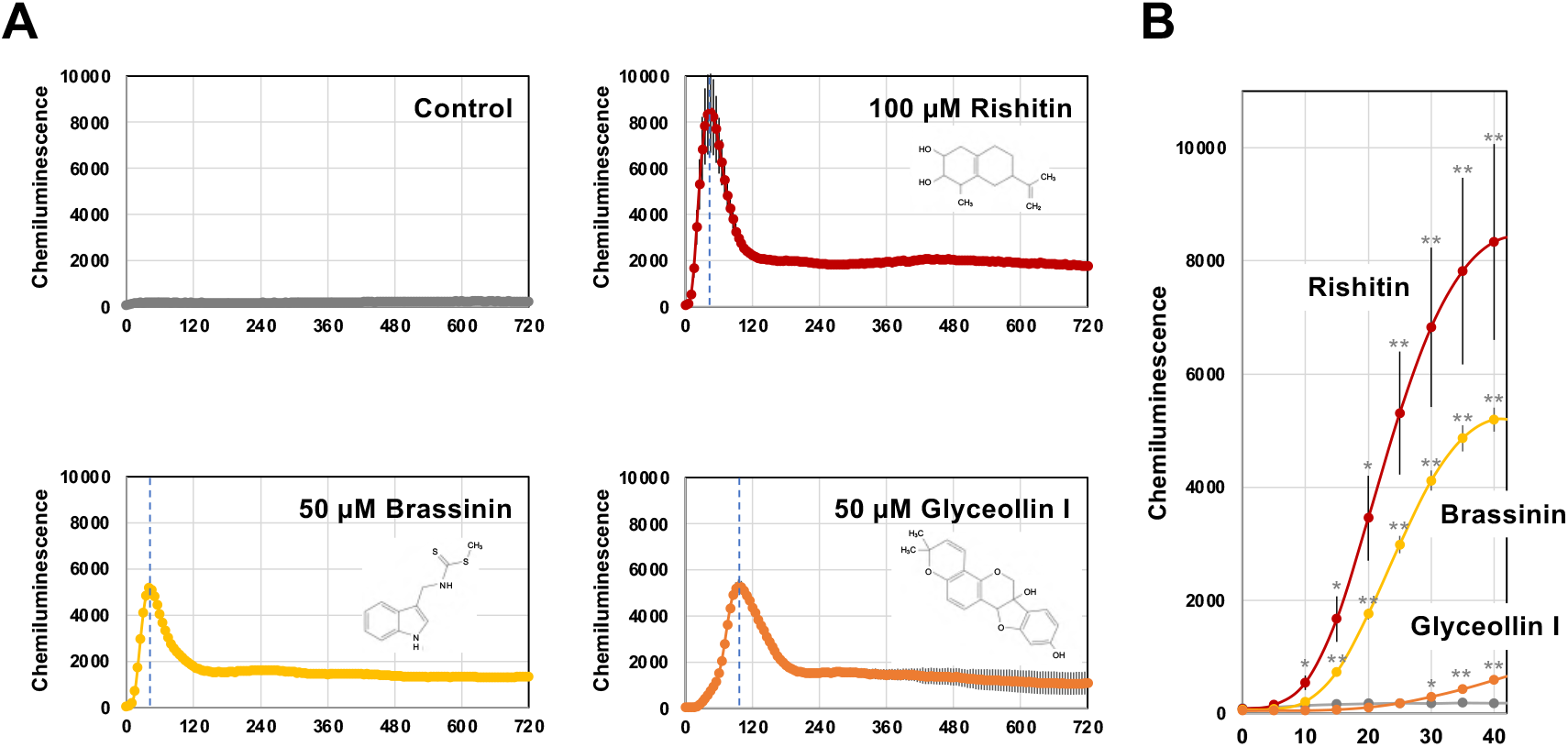
Activation of the *BcatrB* promoter detected in *Botrytis cinerea* transformant expressing *Luciferase* gene under the control of 2 kb *BcatrB* promoter (P_*BcatrB*:*Luc*). **(A)** P_*BcatrB*:*Luc* transformant was incubated in 1% DMSO (Control), 100 µM rishitin, 50 µM brassinin or 50 µM glyceollin containing 50 µM D-Luciferin (substrate of luciferase). Data are mean ± SE (*n* = 3). Dotted vertical lines indicate the time point of highest value for each treatment. **(B)** Activation of *BcatrB* promoter at early time points shown in (A). Asterisks indicate a significant difference from the control as assessed by two-tailed Student’s *t*-test, \*\**P* < 0.01, \**P* < 0.05.

Activation of the *BcatrB* promoter was also tested with brassinin and glyceollin. Treatment of 50 µM brassinin or glyceollin induced transient activation of the *BcatrB* promoter as in the case of rishitin treatment. However, the induction peak of *BcatrB* expression under glyceollin treatment occurred significantly later than those induced by rishitin and brassinin (Figure 7A). Likewise, activation of the *BcatrB* promoter by the brassinin treatment occurred within 15 minutes, whereas weak activation was detected 30 min after glyceollin treatment (Figure 7B). These results suggest that the induction of *BcatrB* expression by rishitin/brassinin and glyceollin might be activated by different regulatory mechanisms.

Previous analysis revealed that some fungicide-resistant *B. cinerea* isolated from fields express high levels of *BcatrB*. A common mutation in these isolates was found in the coding sequence of Zn(II)_2_Cys_6_-type transcription factor mmr1 (Kretschmer et al., 2009), suggesting that developing the transcriptional regulation of gene(s) for drug resistance transporters, like *BcatrB*, is an important process in the evolution of gray mold fungi to become a pleiotropic pathogen. *BcatrB* homologues, and proximal genes/orthologues, are widely conserved in the *Botrytis* genus as well as closely related taxa. This is counter-intuitive to the variability in host ranges driven by phytoalexin tolerance. Thus, it will be interesting to examine whether *Botrytis* sp. with narrow host range can induce the expression of *BcatrB* orthologs in response to phytoalexins from various plant species. The detailed analysis of *BcatrB* promoter may provide clues as to whether there are multiple cis elements involved in finetuning the expression of *BcatrB* across phytoalexin treatments.

## Author contributions

DT designed the research. AB, TK, AA, KI, MO and DT conducted the experiments. AB, MC, AT and DT analyzed data, IS, SC, MO and DT supervised the experiments. AB, MC, IS, SC, and DT contributed to the discussion and interpretation of the results. AB and DT wrote the manuscript. AB, MC and DT edited the manuscript.

## Acknowledgments

We thank Emeritus Prof. Barry Scott (Massey University, New Zealand) for providing *E. festucae* strain Fl1. We also thank Emeritus Prof. David A. Jones (The Australian National University, Australia) for *N. benthamiana* seeds. We are also grateful to Dr. Kenji Asano (National Agricultural Research Center for Hokkaido Region, Japan) and Mr. Yasuki Tahara (Nagoya University, Japan) for providing tubers of potato cultivars. Likewise, we would like to acknowledge the Japanese Ministry of Education, Culture, Sports and Technology (MEXT) and the University of the Philippines Los Banos for allowing Abriel Salaria Bulasag to pursue graduate studies in Japan on scholarship.

