## Supplemental Figs and Tables for "Functional characterization of *Botrytis cinerea* ABC transporter gene *BcatrB* in response to phytoalexins produced in plants belonging to families Solanaceae, Brassicaceae and Fabaceae"

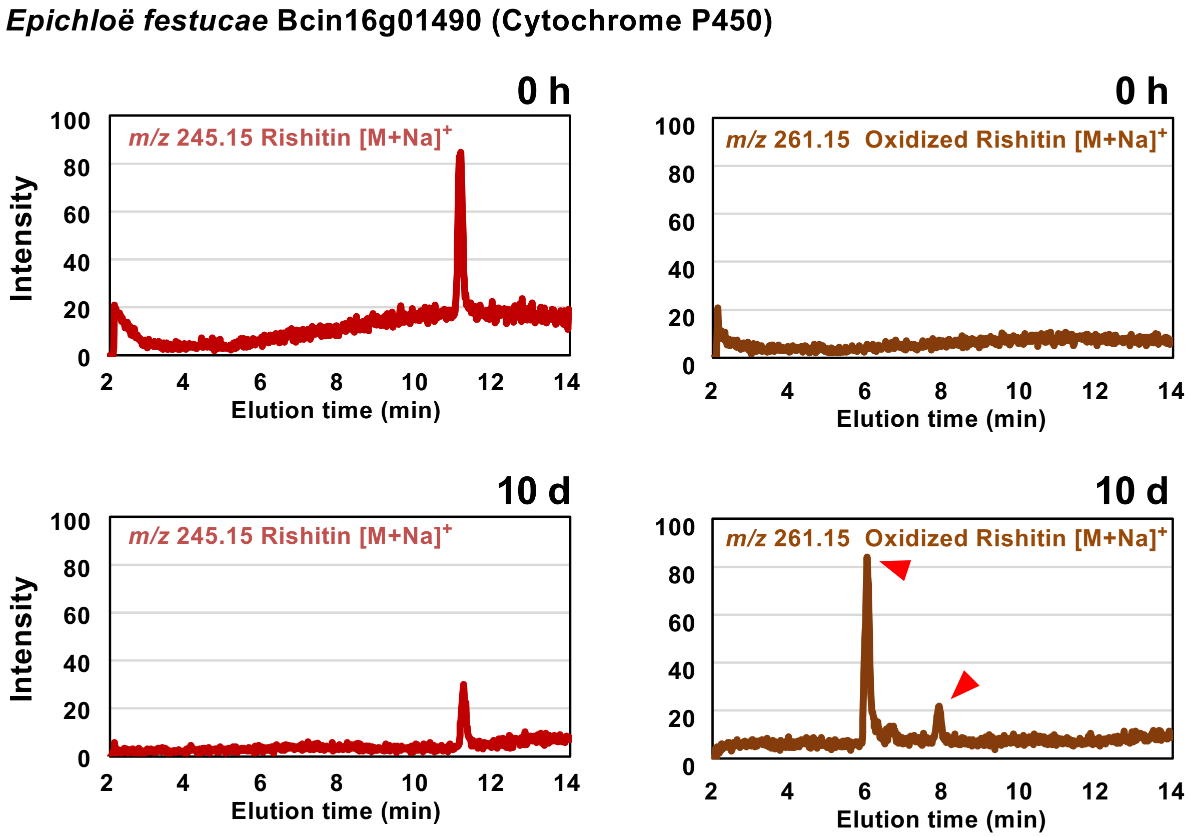

**SUPPLEMENTARY FIGURE S1** | Metabolization of rishitin by *Epichloë festucae* transformants expressing rishitin-induced *B. cinerea* gene Bcin16g01490 encoding cytochrome P450. Mycelial blocks (approx. 1 $mm^3$) of *E. festucae* transformant expressing Bcin16g01490 were incubated in 50 µl of 100 µM rishitin for 0 h or 10 days and rishitin and oxidized rishitin were detected by LC/MS.

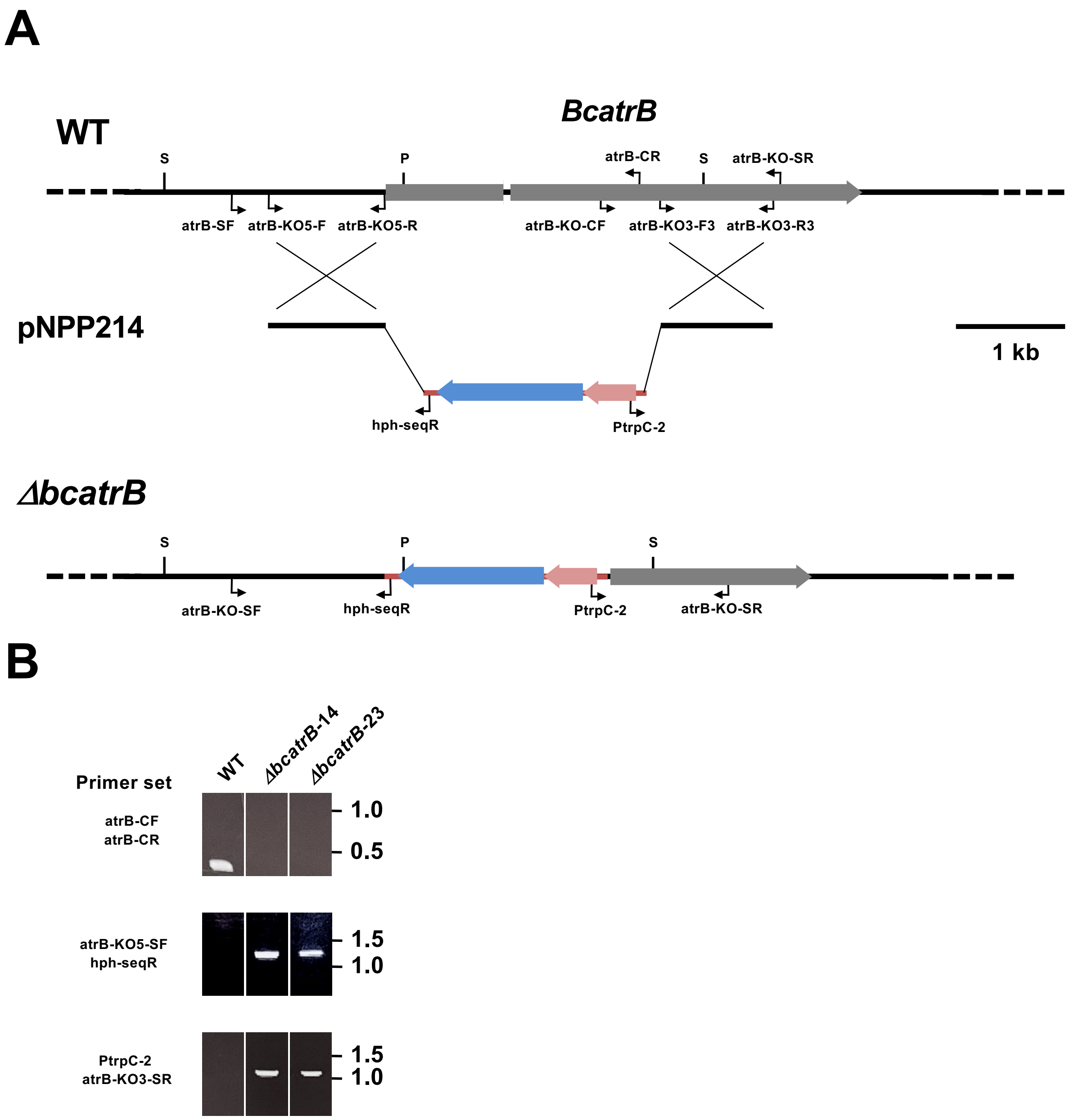

**SUPPLEMENTARY FIGURE S2 |** Targeted gene replacement of the *B. cinerea BcatrB* locus. **(A)** Physical map of the *BcatrB* wild-type (WT) genomic region, linear insert of *BcatrB* replacement construct pNPP214, showing restriction enzyme sites for *Spe*I (S) and *Pst*I (P). The mutated genomic locus of the *BcatrB* deletion mutant (*ΔbcatrB*) is depicted to show homologous recombination of the *hph* cassette. Primers used for the construction of deletion vector and screening for the replacement event are indicated by arrows. **(B)** Confirmation of gene disruption in isolated *ΔbcatrB* strains by PCR. Genomic DNA from *B. cinerea* wild type and *ΔBcatrB* strains were used for PCR with indicated primers.

**Supplementary Table S1.** Fungal strains used in this study.

| Strains | **Relevant characteristics** | **References** |
|---|---|---|
| ***Epichloë festucae*** | | |
| PN2278 (Fl1) | Wild type | Young et al., 2005 |
| Ef-DsRed | | Kayano et al., 2013 |
| Ef-Bcin08g04910 | Fl1/pNPP209; $Hyg^R$ | This study |
| Ef-Bcin16g01490 | Fl1/pNPP208; $Hyg^R$ | This study |
| ***Botrytis cinerea*** | | |
| AI18 | Wild type | Kuroyanagi et al., 2022 |
| *Δ bcatrB* -14–2 | Fl1/pNPP214; $Hyg^R$ | This study |
| *Δ bcatrB* -23-3 | Fl1/pNPP214; $Hyg^R$ | This study |
| *P_BcatrB* :*GFP* | Fl1/pNPP213; $Hyg^R$ | This study |
| *P_BcatrB* :*Luc* | Fl1/pNPP212; $Hyg^R$ | This study |

**Supplementary Table S2.** Plasmids used in this study

| Vector name | Base vector | | Insert | Primers used to amplify insert | References | note |
|---|---|---|---|---|---|---|
| **Base vectors** | | | | | | |
| pPN94 | – | – | – | – | Takemoto et al. 2006 | Base vector for gene expression under TEF promoter, $Amp^R/Hyg^R$ |
| pNPP150 | – | – | – | – | Niones and Takemoto 2015 | Base vector for gene knockout (*HSVtk* marker), $Amp^R/Hyg^R$ |
| pNPP170-BcGFP | – | – | – | – | Kuroyanagi et al. 2022 | Base vector for the promoter analysis using BcGFP marker, $Amp^R/Hyg^R$ |
| pNPP210 | – | – | – | – | Kuroyanagi et al. 2022 | Base vector for promoter analysis using Luciferase marker, $Amp^R/Hyg^R$ |
| **Plasmids for gene expression in *E. festucae* and *B. cinerea*** | | | | | | |
| pNPP208 (pPN94-Bcin16g01490) | pPN94 | *Bam* HI/ *Not* I | Bcin16g01490 | IF-94-16g01490-F, IF-94-16g01490-R | This study | Constitutive expression of Bcin16g01490 |
| pNPP209 (pPN94-Bcin08g04910) | pPN94 | *Bam* HI/ *Not* I | Bcin08g04910 | IF-94-08g04910-OR-F, IF-94-08g04910-OR-R | This study | Constitutive expression of Bcin08g04910 |
| pNPP213 (pNPP170-P_Bcin08g00930-BcGFP) | pNPP170-BcGFP | *Sac* I/ *Sma* I | *BcatrB* promoter (1 kb) | pNPP170-P_8g00930-F, pNPP170-P_8g00930-R | This study | Expression of GFP under the control of 1 kb *BcatrB* promoter |
| pNPP214 (pNPP150-BcatrB-KO) | pNPP150 | *Sal* I/ EcoRI | 5'*BcatrB* -PtrpC-hph-3'*BcatrB* | atrB-KO5-F, atrB-KO5-R, atrB-KO3-F, atrB-KO3-R | This study | Knockout vector for *BcatrB* |
| pNPP212 (P_*BcatrB* :*Luc* 2kb) | pNPP210 | *Eco* RI | *BcatrB* promoter (2kb) | pNPP210-P_BcatrB_2kb, pNPP210-P_13g00710-R | This study | Expression of Luc under the control of 2 kb *BcatrB* promoter |

**Supplemnetary Table S3.** Primers for sequencing, vector construction used in this study.

| Primer name | Sequence 5'→3' |
|---|---|
| **Primers for sequencing of vectors** | |
| Ptef-seq | TAACCTCTCTTCAGAAAG |
| TtrpC-seq | TCTGGAAGAGGTAAACCCG |
| pII99-3 | GGCTGGCTTAACTATGCG |
| PtrpC-2 | CAAATTTTGTGCTCACCG |
| pII99-2 | CGGTATCAGCTCACTCAA |
| TtrpC-1 | ACACACATTCATCGTAGG |
| **Primers for construction of expression and knock out vectors** | |
| IF-94-16g01490-F | AACCTCTAGAGGATCATGTCGCCAGCACTCTTCGA |
| IF-94-16g01490-R | ACGTTAAGTGCGGCCTTAAACTCTTCTCTTGATCT |
| IF-94-8g04910-OR-F | AACCTCTAGAGGATCATGAGTGTCATCAAAAACGT |
| IF-94-8g04910-OR-R | ACGTTAAGTGCGGCCTTACTTGACATACTTCTTCA |
| atrB-KO5-F | ATGCCTGCAGGTCGACCTAGCTACGATTGATGGAA |
| atrB-KO5-R | ATCCTCTAGAGTCGAGATGGCAATTGAAGTATTGA |
| atrB-KO3-F3 | TACCGAGCTCGAATTTGTACAAGGGTGGGTTAAGC |
| atrB-KO3-R3 | TATCATCGATGAATTCCGTGAAGAGACGTAATTGC |
| pNPP170-P_13g00710-F | CCAAGCTGGGTACCGTGAGAAACAAAGGTAAGACA |
| pNPP170-P_13g00710-R | AACCATGGTGAACCCGATGGCAATTGAAGTATTGA |
| pNPP210-P_13g00710-R | CGTCCTCCATGAATTGATGGCAATTGAAGTATTGA |
| pNPP210-P_BcatrB_2kb | TACCGAGCTCGAATTTCACCGAAACTTCAGCCTTC |
| **Primers for genomic PCR** | |
| atrB-KO5-SF | TCGGGCGAGTGCAGCATTTC |
| atrB-KO3-3-SR | AGGTTGCAGTTGAGCCATGA |
| atrB-CF | GCCTATGGCTTTTCGGCTAT |
| atrB-CR | CAGCCTTGAGAATAGCAGTA |

Extension sequences for In-fusion reaction are in red letters.

**Supplementary Table S4.** Expression of GFP in *Botrytis cinerea* P_*BcatrB*:*GFP* transformant in different plant species.

| Family | Host plant | Common name | Disease symptom | Expression of GFP in *B. cinerea* P_*BcatrB*:*GFP* |
|---|---|---|---|---|
| Solanaceae | *Nicotiana benthamiana* | Benth | + | + |
| Solanaceae | *Solunum lycopercicum* | Tomato | + | +* |
| Solanaceae | *Solunum melongena* | Eggplant | + | + |
| Solanaceae | *Capsicum annuum* | Bell pepper | + | - |
| Brassicaceae | *Brassica oleracea* var. *capitata* | Cabbage | + | + |
| Brassicaceae | *Brassica oleracea* var. *italica* | Broccoli | + | + |
| Brassicaceae | *Arabidopsis thaliana* | Thale cress | + | + |
| Fabaceae | *Trifolium pratense* | Red clover | + | +++ |
| Fabaceae | *Trifolium repens* | White clover | + | ++ |
| Fabaceae | *Glycine max* | Soybean | + | ++ |
| Fabaceae | *Arachis hypogaea* | Peanut | + | +++ |
| Fabaceae | *Phaseolus vulgaris* | Green bean | + | +++ |
| Cucurbitaceae | *Cucurbita maxima* | Winter squash | + | - |
| Cucurbitaceae | *Cucumis sativus L.* | Cucumber | + | - |
| Cucurbitaceae | *Citrullus lanatus* | Watermelon | + | - |
| Vitaceae | *Vitis × labruscana* | Grape | + | +* |
| Vitaceae | *Parthenocissus tricuspidata* | Boston Ivy | + | - |
| Lamiaceae | *Ocimum basilicum* | Basil | + | - |
| Rosaceae | *Cerasus × yedoensis* | Sakura | + | - |
| Hydrangeaceae | *Hydrangea macrophylla* | Hydrangea | + | - |
| Caryophyllaceae | *Dianthus caryophyllus* | Carnation | + | - |
| Asteraceae | *Taraxacum officinale* | Dandelion | + | - |
| Geraniaceae | *Geranium caroliniaum* | Geranium | + | - |
| Amaryllidaceae | *Allium fistulosum* | Welsh onion | + | - |
| Oxalidaceae | *Oxalis corniculata* | Creeping woodsorrel | + | - |

* GFP expression in fruits but not in leaves.
